# Replication of SARS-CoV-2 in human respiratory epithelium

**DOI:** 10.1101/2020.03.20.999029

**Authors:** Aleksandra Milewska, Anna Kula-Pacurar, Jakub Wadas, Agnieszka Suder, Artur Szczepanski, Agnieszka Dabrowska, Katarzyna Owczarek, Marek Ochman, Tomasz Stacel, Zenon Rajfur, Pawel Labaj, Wojciech Branicki, Krzysztof Pyrc

## Abstract

SARS-CoV-2 emerged by the end of 2019 to rapidly spread in 2020. At present, it is of utmost importance to understand the virus biology and to rapidly assess the potential of existing drugs and develop new active compounds. While some animal models for such studies are under development, most of the research is carried out in the Vero E6 cells. Here, we propose fully differentiated human airway epithelium cultures as a model for studies on the SARS-CoV-2. Further, we also provide basic characteristics of the system.

## Introduction

Coronaviruses constitute a large family of RNA viruses that infect mainly mammals and birds. In humans, there are four species associated with mild-to-moderate respiratory infections. While these viruses are present in the human population for a long time, they are believed to enter the human population in a zoonotic event, and one may speculate that they may have caused epidemics similar to the one observed for the SARS-CoV-2. Time to the most recent ancestor analysis suggests that human coronavirus HCoV-NL63 is the oldest species in humans, followed by its cousin HCoV-229E and two betacoronaviruses, which emerged in humans in a relatively near past^1,2, 3,4^. In the 21^st^ century, we already faced the emergence of the three novel coronaviruses in humans, of which SARS-CoV disappeared after one season never to come back, and MERS-CoV never fully crossed the species border, as its transmission between humans is highly ineffective^5,6,7^. The 2019 zoonotic transmission, however, resulted in the emergence of a novel human coronavirus, which seems to carry an optimal set of features allowing for its rapid spread with considerable mortality. Whether the virus will become endemic in humans is an open question^8,9,10^.

At present, the studies on the virus are carried out using a surrogate system based on the immortalized simian Vero E6 cell line^11^. While this model is convenient for diagnostics and testing of some antiviral drugs, it has serious limitations and does not allow for the understanding of virus biology and evolution. To make an example, the entry route of human coronaviruses varies between the cell lines and differentiated tissue, not mentioning the immune responses or virus-host interactions^12,13,14^.

Here we used the fully differentiated epithelium cultures to study the infection with the novel human coronavirus SARS-CoV-2. We observed an efficient replication of the virus in the tissue, with the maximal replication at 2 days post-infection. At the time of the study no antibodies were available. Therefore we developed immuno-FISH to show that the virus infects primarily ciliated cells of the respiratory epithelium.

## Results and discussion

The HAE cultures reconstitute the tissue lining the conductive airways of humans. Fully differentiated, are among the best tools for studying the viral infection in a natural microenvironment^15^. These air-liquid interphase cultures contain a number of cell types (e.g., basal, ciliated, and goblet). At the same time, they also functionally reflect the natural tissue with extensive crosstalk and production of protective mucus and surfactant proteins^16,17,18^. The cultures were previously shown by us and others to be superior to the standard cell lines in terms of ability to support coronaviral replication of the HCoV-HKU1, but also as a model to study the biology of the infection^19^. To make an example, human coronaviruses were shown some time ago to use a very different entry pathway in immortalized cell lines and in the natural human epithelium. While in the first one they enter via pH-dependent endocytic pathway, in the latter one they utilize surface serine proteases as TMPRSS2 or kallikreins for activation and the fusion occurs on the cell surface. This may have grave consequences not only for the basic science, but also the antiviral drug development^12, 13, 14, 20^.

Here we verified whether HAE cultures may be used to study the SARS-CoV-2 infection and identified the cellular targets in the tissue. First, the HAE cultures were inoculated with the SARS-CoV-2 stock and cultured for 5 days. Every day (days 0-4) the apical and basolateral release of the virus was evaluated with the RT-qPCR and the results for the apical release of the virus are presented in **Figure 1**.

**Figure 1.**
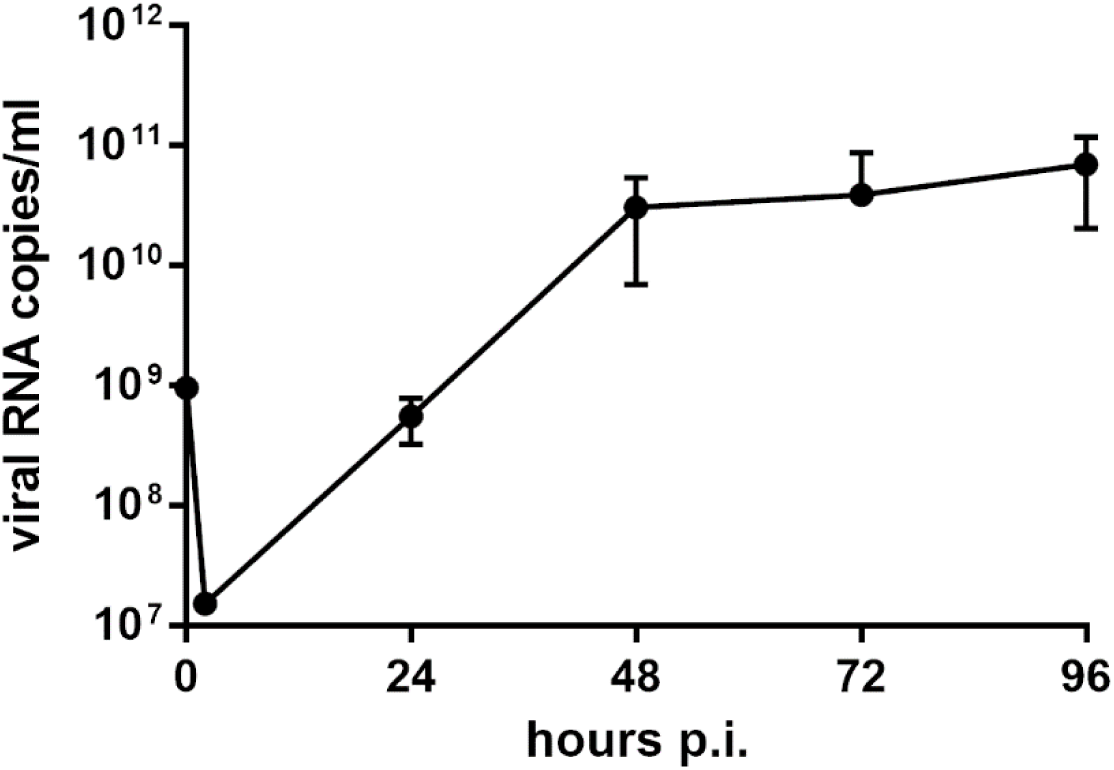
SARS-CoV-2 replicates in HAE cultures. Replication of SARS-CoV-2 was evaluated using an RT-qPCR, and the data are presented as RNA copy number per ml. The experiment was carried out in triplicate, and average values with standard deviation are presented.

Clearly, the increase in virus titer on the apical side is visible already 24 h post-inoculation, to reach the plateau at 48 h post-inoculation. We did not observe any release of the virus from the basolateral side of the HAE culture and therefore we do not show the relevant data on the graph. The results we observe are consistent with the previously reported polarity of the HAE cultures and apical infection / apical release reported previously for other human coronaviruses. Similarly as for other human coronaviruses, the apical-apical polarity of SARS-CoV-2 infection-release restricts the virus to the airway lumen^16^.

Further, we sampled the tissue at 96 h post-infection, to verify whether the subgenomic mRNAs are present. The analysis was carried out with RT-PCR and the results are presented in **Figure 2**.

**Figure 2.**
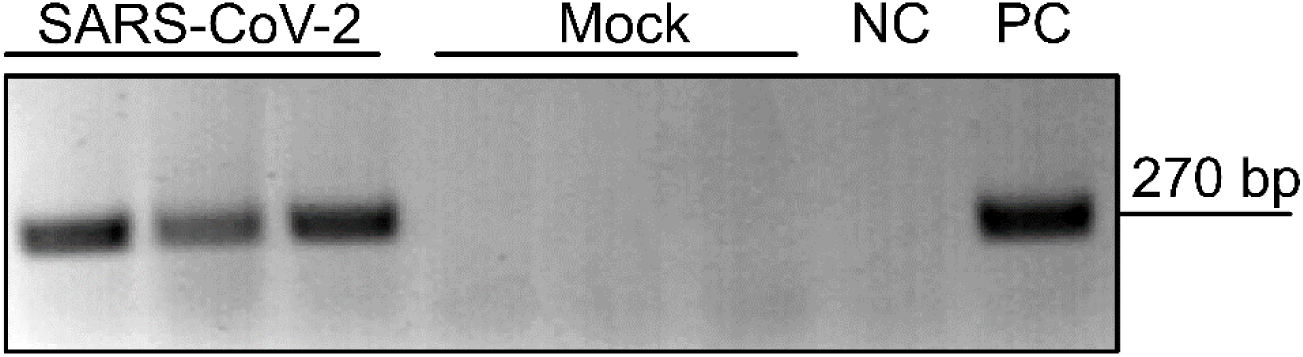
sg mRNAs of the SARS-CoV-2 in HAE cultures. The presence of the N sg mRNAs 4 days p.i. in the HAE cultures infected with the SARS-CoV-2 was evaluated using an RT-PCR. NC = negative control, PC = positive control.

The analysis clearly showed that the sg mRNA are abundant in the infected HAE cultures. As this is generally considered to be the hallmark of an active replication, we believe that it provides sufficient proof that the virus is indeed actively replicating in the cultures.

Next, we made an effort to visualize the infection in the tissue. As at the time of the study no antibody for the confocal microscopy was available, we developed an immuno-FISH assay, where the viral RNA was visualized in the context of the cell using 20 sequence-specific probes and signal amplification. At the same time, the β-tubulin was visualized using specific antibodies to visualize the ciliated cells. Obtained results are shown in **Figure 3**.

**Figure 3.**
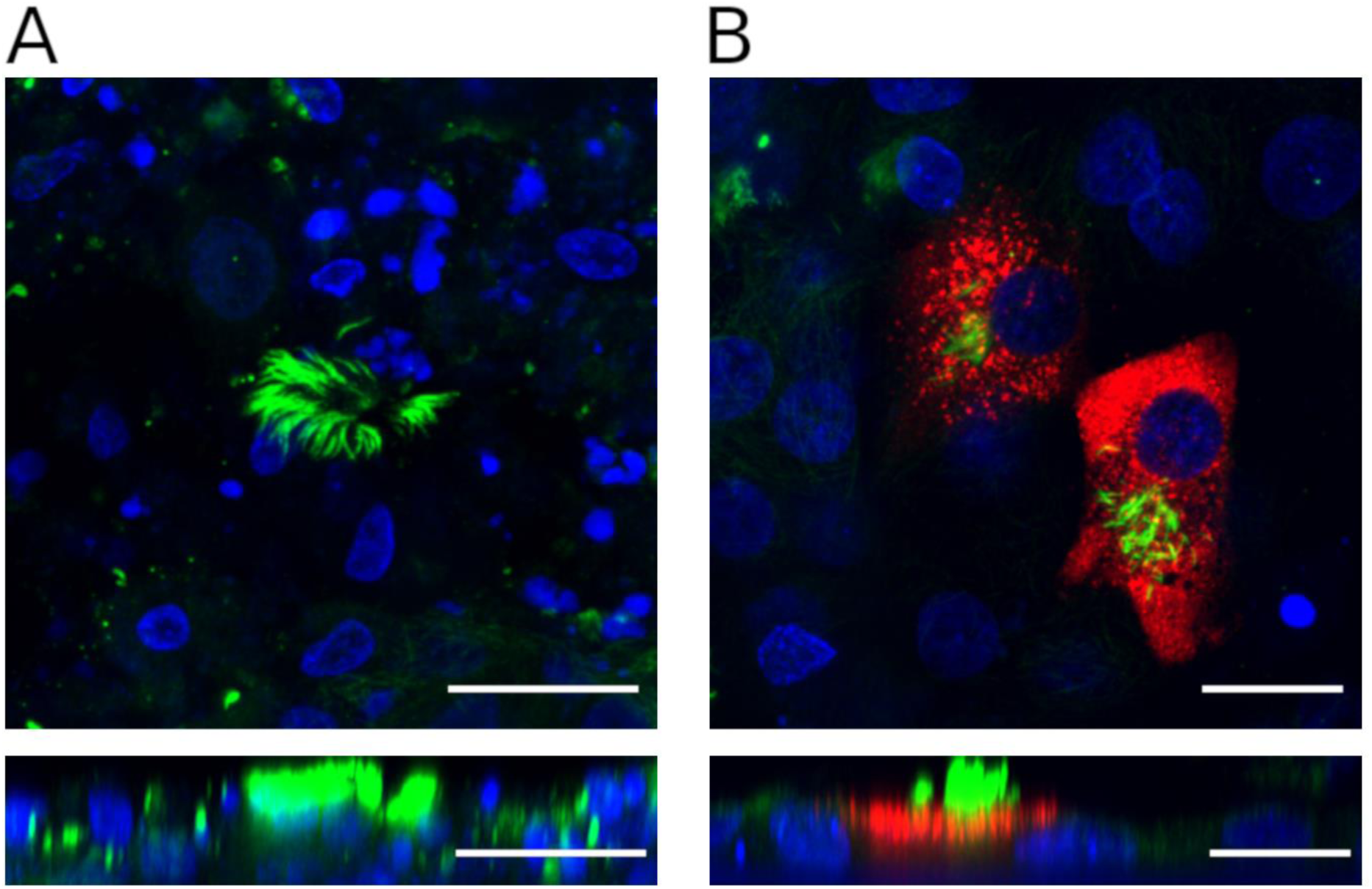
SARS-CoV-2 infects ciliated cells of the human airway epithelium. Three-dimensional Immuno-RNA FISH demonstrating localization of SARS-CoV-2 subgenomic RNA in ciliated HAE cultures. Three-dimensionally reconstructed confocal image stacks of cells infected with SARS-CoV-2 (A) and mock control cells (B). The bottom lanes of panels A and B show the xz plane in orthogonal views. SARS-CoV-2 RNA is visualized by FISH using a set of probes against viral nucleocapsid RNA and is shown in red. Cilia are visualized by an anti-β5 tubulin antibody and are shown in green. Nuclei are stained with DAPI and are shown in blue. Bar = 20µM.

Summarizing, we show that the SARS-CoV-2 effectively replicates in the HAE cultures and that this *ex vivo* model constitutes a convenient tool to study the viral infection. We also show that the virus infects ciliated cells. The infection is polarized - the infection and release occurs at the apical side of the epithelium. It is worth to note that in the absence of the immunodetection tools the new generation of immune-FISH tools offers an interesting alternative.

## Materials and Methods

### Cell culture

Vero E6 (*Cercopithecus aethiops*; kidney epithelial; ATCC: CRL-1586) cells were maintained in DMEM (Thermo Fisher Scientific, Poland) supplemented with 3% FBS (heat-inactivated fetal bovine serum; Thermo Fisher Scientific, Poland) and streptomycin (100 μg/ml), penicillin (100 U/ml), and ciprofloxacin (5 μg/ml). Cells were cultured at 37°C in atmosphere containing 5% CO_2_.

### Human airway epithelium (HAE) cultures

Human epithelial cells were isolated from conductive airways resected from transplant patients. The study was approved by the Bioethical Committee of the Medical University of Silesia in Katowice, Poland (approval no: KNW/0022/KB1/17/10 dated 16.02.2010). Written consent was obtained from all patients. Cells were mechanically detached from the tissue after protease treatment and cultured on plastic in BEGM media. Subsequently, cells were transferred onto permeable Transwell insert supports (ϕ = 6.5 mm) and cultured in BEGM media. After the cells reached full confluency, the apical medium was removed, and the basolateral medium was changed to ALI. Cells were cultured for 4-6 weeks to form differentiated, pseudostratified mucociliary epithelium. All experiments were performed in accordance with relevant guidelines and regulations.

### Virus

SARS-CoV-2 (isolate 026V-03883; kindly granted by Christian Drosten, Charité – Universitätsmedizin Berlin, Germany and provided by the European Virus Archive - Global (EVAg); https://www.european-virus-archive.com/). Virus stock was prepared by infecting fully confluent Vero E6 cells at a TCID_50_ of 400 per ml. Three days after inoculation, supernatant from the cultures was aliquoted and stored at −80°C. Control Vero E6 cell supernatant from mock-infected cells was prepared in the same manner. Virus yield was assessed by titration on fully confluent Vero E6 cells in 96-well plates, according to the method of Reed and Muench. Plates were incubated at 37°C for 2 days, and the cytopathic effect (CPE) was scored by observation under an inverted microscope.

### Virus infection

Fully differentiated human airway epithelium (HAE) cultures were inoculated with the SARS-CoV-2 at a TCID_50_ of 1000 per ml (as determined on Vero E6 cells). Following 2 h incubation at 37°C, unbound virions were removed by washing with 200 μl of 1 × PBS, and HAE cultures were maintained at an air-liquid interphase for the rest of the experiment. To analyze virus replication kinetics, each day p.i., 100 μl of 1 × PBS was applied at the apical surface of HAE and collected following the 10 min incubation at 32°C. All samples were stored at −80°C and analyzed using RT-qPCR.

Additionally, 48 h post-infection, selected HAE cultures were collected, and the presence of sg mRNA was determined as hallmarks of an active infection.

### Isolation of nucleic acids and reverse transcription (RT)

Viral DNA/RNA Kit (A&A Biotechnology, Poland) was used for nucleic acid isolation from cell culture supernatants and Fenozol (A&A biotechnology, Poland) was used for total RNA isolation from cells. RNA was isolated according to the manufacturer’s instructions. cDNA samples were prepared with a High Capacity cDNA Reverse Transcription Kit (Thermo Fisher Scientific, Poland), according to the manufacturer’s instructions.

### Quantitative PCR (qPCR)

Viral RNA was quantified using qPCR (CFX96 Touch Real-Time PCR Detection System, Bio-Rad, Poland). cDNA was amplified using 1 × qPCR Master Mix (A&A Biotechnology, Poland), in the presence of probe (100 nM, FAM / BHQ1, ACT TCC TCA AGG AAC AAC ATT GCC A) and primers (450 nM each, CAC ATT GGC ACC CGC AAT C and GAG GAA CGA GAA GAG GCT TG). The heating scheme was as follows: 2 min at 50°C and 10 min at 92°C, followed by 30 cycles of 15 s at 92°C and 1 min at 60°C. In order to assess the copy number for N gene, standards were prepared and serially diluted.

### Detection of SARS-CoV-2 N sg mRNA

Total nucleic acids were isolated from virus or mock-infected cells at 4 days p.i. using Fenozol reagent (A&A Biotechnology, Poland), according to the manufacturer’s instructions. Reverse transcription was performed using a high-capacity cDNA reverse transcription kit (Life Technologies, Poland), according to the manufacturer’s instructions. Viral cDNA was amplified in a 20 µl reaction mixture containing 1 × Dream *Taq* Green PCR master mix (Thermo Fisher Scientific, Poland), and primers (500 nM each). The following primers were used to amplify SARS-CoV-2 subgenomic mRNA (sg mRNA): common sense primer (leader sequence), 5 – TAT ACC TTC CCA GGT AAC AAA CCA -3’; nucleocapsid antisense, 5’ – GTA GCT CTT CGG TAG TAG CCA AT – 3’. The conditions were as follows: 3 min at 95°C, 35 cycles of 30 s at 95°C, 30 s at 52°C, and 20 s at 72°C, followed by 5 min at 72°C and 10 min at 4°C. The PCR products were run on 1% agarose gels (1Tris-acetate EDTA [TAE] buffer) and analyzed using molecular imaging software (Thermo Fisher Scientific, Poland).

### RNA Fluorescent *in situ* Hybridization (RNA-FISH) and Immunofluorescence

HAE cultures were infected with SARS-CoV-2 [TCID_50_=1000, as assessed for the Vero E6 cells] and fixed at 5 days post-infection with 3.7% paraformaldehyde (PFA) overnight. The next day, cells were subjected to RNA-FISH protocol using hybridization chain reaction (HCR) technology from Molecular Instruments, Inc. Briefly, cells were permeabilized with 100% methanol overnight and then subjected to grated rehydration with methanol/PBS, Tween 0.1%. The set of DNA HCR v3.0 probes complementary to SARS-CoV-2 nucleocapsid RNA was incubated for 12 h at 37°C, extensively washed, and hybridized with HCR amplifiers for 12 h at room temperature in the dark. Next, cells were subjected to immunostaining with antibodies against mouse β5-tubulin from Santa Cruz Biotechnology (sc-134234, 1:100), rinsed three times with PBS, 0.1% Tween-20 and followed by 1 h incubation with Alexa fluorophore 488 secondary antibodies (Invitrogen, 1:400). The cells were finally washed three times with PBS, 0.1% Tween-20, cell nuclei were stained with DAPI (4 =,6 =-diamidino-2-phenylindole) (Thermo Fisher Scientific, D1306) and mounted on slides with Prolong diamond antifade mounting medium (Invitrogen, P36970). Fluorescent images were acquired using a Zeiss LSM 710 confocal microscope (Carl Zeiss Microscopy GmbH) with ZEN 2012 SP1 Black Edition and processed in ImageJ Fiji (National Institutes of Health, Bethesda, MD, USA).

## Funding

This work was supported by the funds provided by the Ministry of Science and Higher Education for research on the SARS-CoV-2 and grants from the National Science Center (grants UMO-2017/27/B/NZ6/02488 to KP and UMO-2018/30/E/NZ1/00874 to AKP).

## Conflicts of Interest

The authors declare no conflict of interest. The funders had no role in the design of the study; in the collection, analyses, or interpretation of data; in the writing of the manuscript, or in the decision to publish the results.

